# Deep White-Matter Pathways Mediate the Link Between Docosahexaenoic Acid (DHA) Status and Cognitive Performance in Adolescence

**DOI:** 10.1101/2025.11.20.689543

**Authors:** Alejandra Figueroa-Vargas, Rodrigo Valenzuela, Patricia Soto-Icaza, Francisco Zamorano, Cristián Larraín, Claudio Silva, Pamela Guevara, Sebastian Navarrete, Atilio Almagià, Violeta Arancibia, Cynthia Barrera, Daniza Ivanovic, Pablo Billeke

## Abstract

Docosahexaenoic acid (DHA) is a polyunsaturated fatty acid enriched in neuronal membranes and myelin and associated with cognitive performance. However, nutritional interventions show inconsistent cognitive effects, partly due to limited knowledge of the neural pathways linking DHA status to human cognition during sensitive periods of white-matter maturation, such as adolescence. We addressed this gap by studying 99 adolescents drawn from both extremes of performance on a national scholastic examination. Participants completed assessments of scholastic achievement (SA) and intellectual ability (IA), provided erythrocyte DHA samples, and underwent multimodal MRI, including diffusion, T1-weighted, and T2-weighted imaging. Independent component analysis and Bayesian multivariate LASSO models identified brain components jointly associated with DHA and cognition. Across four MRI modalities, a single deep white-matter component consistently emerged as the strongest shared pathway linking DHA with cognition. Tract-resolved analyses highlighted predominant contributions from the fornix and thalamus–temporal fasciculus, with additional subcortical and cortical involvement. In joint models, these components predicted SA and IA after accounting for DHA and other fatty acids, consistent with an indirect, mediation-like pathway. These findings move beyond DHA–behavior correlations by identifying specific neuroanatomical pathways through which a modifiable dietary factor relates to adolescent learning and intellectual performance, offering mechanistic insight relevant to neuroscience, nutrition, and education.

**Significance Statement:** Adolescence is a sensitive period for the maturation of white-matter pathways that support learning and reasoning. Docosahexaenoic acid (DHA), an essential dietary fatty acid enriched in neuronal membranes and myelin, has been linked to cognitive performance, yet the neural mechanisms underlying this association remain unclear. Using multimodal MRI and Bayesian multivariate modeling in adolescents with high or low scholastic performance, identify a specific deep white-matter pathway—centered on the fornix and the thalamus–temporal fasciculus—as the principal route connecting DHA status with scholastic achievement and intellectual ability. Additional subcortical and cortical contributions reveal a coordinated system-level architecture. These findings move beyond correlations by providing mechanistic insight into how a modifiable nutritional factor relates to cognitive development, with implications for neuroscience, public health, and education.

## Introduction

The emergence of the cognitive abilities required to meet the demands of modern societies arises from complex, multifactorial biological processes. Among these, nutrition has been historically underestimated despite its profound influence on brain network organization [1]. Docosahexaenoic acid (DHA; C22:6n-3), a major n-3 polyunsaturated fatty acid (n-3 PUFA), is essential for maintaining neuronal membrane structure, synaptic function, and myelin integrity [2,3]. DHA accumulates rapidly during late fetal development and early childhood, supporting synaptogenesis, membrane fluidity, and neuroprotection across the lifespan [4]. Because endogenous DHA synthesis from α-linolenic acid (ALA; C18:3n-3) is highly limited, adequate levels depend on dietary intake [5]. Although DHA represents a modifiable nutritional factor with clear biological relevance, dietary interventions alone often yield inconsistent cognitive benefits [6,7], highlighting a crucial mechanistic gap: how do specific nutritional factors shape identifiable neural pathways that support cognitive performance?

Adolescence—spanning ages 10 to 24 years [8]—represents a sensitive developmental period in which nutritional factors may play a particularly influential role. Evidence suggests that higher erythrocyte DHA is associated with better attentional performance in typically developing adolescents [9], whereas different n-3 PUFAs, such as eicosapentaenoic acid (EPA), may show dissociable associations with memory in clinical populations [10]. These findings underscore the need to distinguish the roles of specific PUFAs in adolescent cognition. Moreover, while dietary DHA correlates weakly with cognitive performance [11], erythrocyte DHA provides a more robust biomarker—reflecting long-term intake, correlating strongly with brain DHA in animal models [12], and showing more consistent associations with cognitive and behavioral outcomes in children [13–15].

Scholastic achievement (SA) and intellectual ability (IA) capture complementary dimensions of cognitive functioning. IA reflects controlled, laboratory-based assessments of reasoning, whereas SA reflects the expression of these abilities in real-world, socially embedded educational contexts [16]. Together, they offer a comprehensive window into individual cognitive potential and its realization. Nutritional biomarkers such as DHA provide an important biological layer through which to understand interindividual variability. Higher DHA has been linked not only to improved cognitive performance—including memory, attention, executive function, and processing speed—but also to lower suicide risk [17,18]. Although DHA may modulate neuroinflammatory pathways, as suggested by TSPO PET imaging [19], these effects have not been consistently tied to cognitive outcomes, pointing instead toward structural pathways as likely mediators of DHA-related cognitive differences.

Despite evidence associating DHA with cognition, the neurobiological pathways mediating this relationship remain poorly understood—particularly during adolescence, when myelination, synaptic pruning, and large-scale network refinement proceed rapidly [20–22]. Prior work in older adults suggests that nutritional status may influence cognitive performance indirectly through structural and functional connectivity [23–26], but analogous evidence in adolescents is scarce. During this period, white-matter tracts mature in a back-to-front pattern [20], cortical myelination advances along protracted, region-specific trajectories [21,27,28], and subcortical structures—including the thalamus, pallidum, putamen, and hippocampus—continue to undergo dynamic changes linked to learning, memory, and executive function [29–33].

Taken together, current evidence suggests that brain systems may serve as key mediators linking DHA status to cognitive development. However, no study has yet integrated multimodal MRI features with behavioral measures of both academic achievement and reasoning ability to identify system-level neural pathways through which DHA relates to adolescent cognition. Given DHA’s established roles in membrane fluidity, synaptogenesis, myelination, and neuroprotection [34,35], we hypothesized that distinct patterns of brain organization—including white-matter integrity, subcortical morphology, cortical structure, and cortical myelin content—would mediate the association between DHA status and SA/IA. Using a multivariate, multimodal framework, this study seeks to uncover the coordinated neural mechanisms through which nutritional status shapes cognitive development during adolescence—a period uniquely sensitive to both neurobiological plasticity and environmental influences.

## Results

### Sample

As previously described [36–39], participants were drawn from a representative cohort of school-age students who had completed the University Selection Test (PSU), selected by low and high SA [36,38,39]. A total of 99 participants were successfully scanned after the PSU (see *Methods* for details). Participants’ age ranged from 17.3 y to 20.3 y (mean age = 18.2 ± 0.5 y). Menarcheal age did not differ significantly between females in the high SA (12.6 ± 1.2) and low SA groups (12.5 ± 1.2; F = 0.12; p = 0.7359).

### Socioeconomic status (SES), Scholastic Achievement, Intellectual Ability, Prenatal, postnatal, and current nutritional status

As previously reported [37], SES distribution differed significantly by sex and group (p < 0.0001). High SA participants were predominantly from high/medium SES backgrounds, whereas most Low SA females belonged to low SES categories. SES and related indicators correlated positively with PSU performance in both language and mathematics (p < 0.0001). Intelligence assessment scores (Raven’s Progressive Matrices) differed significantly across groups (p < 0.0001), with High SA participants achieving higher scores. BMI did not differ significantly between groups. Additional details have been described previously [37].

### White-matter integrity, DHA, and Scholastic Achievement, and Intellectual Ability

Prior work shows that circulating DHA relates to cognitive performance, including SA and IA, an association also observed in the original cohort from which the present subsample was drawn [38]. We therefore asked whether white-matter microstructure mediates part of the DHA cognitive performance relationship. We quantified erythrocyte DHA in the neuroimaging subsample and derived fractional anisotropy of white matter (WM-FA) from diffusion-weighted MRI across 36 long-range tracts. To reduce dimensionality, we applied independent component analysis (ICA) to the tract-level WM-FA metrics and retained the set of components explaining 90% of the variance. We then modeled the links among DHA, WM-FA components, IA, and SA using Bayesian multivariate Least Absolute Shrinkage and Selection Operator (Lasso).

As an initial screen, we fit univariate Bayesian Lasso models, testing each WM–FA independent component separately for associations with DHA, SA, and IA, controlling by sex and SES. This analysis identified a single component (IC5) showing convergent evidence: the same component was associated with higher DHA, better SA, and higher IA, all in directionally consistent ways. Specifically, DHA ∼ FA (IC5): posterior median = –0.20, 95% HDI = [–0.30, –0.04], p_LASSO_MCMC_ = 0.009; SA ∼ FA (IC5): posterior median = –0.18, 95% HDI = [–0.30, –0.02], p_LASSO_MCMC_ = 0.010; IA ∼ FA (IC5): posterior median = –0.24, 95% HDI = [–0.40, –0.08], p_LASSO_MCMC_ = 0.002. Although IC5 showed a negative association at the component level, further inspection revealed the underlying positive contributions of specific tracts. By multiplying the posterior distribution of the IC5 coefficient by the tract loadings that constitute this component, we observed positive associations for the FA of the bilateral fornix and the bilateral thalamus–temporal fasciculus, indicating that these tracts primarily drive the observed relationships (See Figure 1).

**Figure 1.**
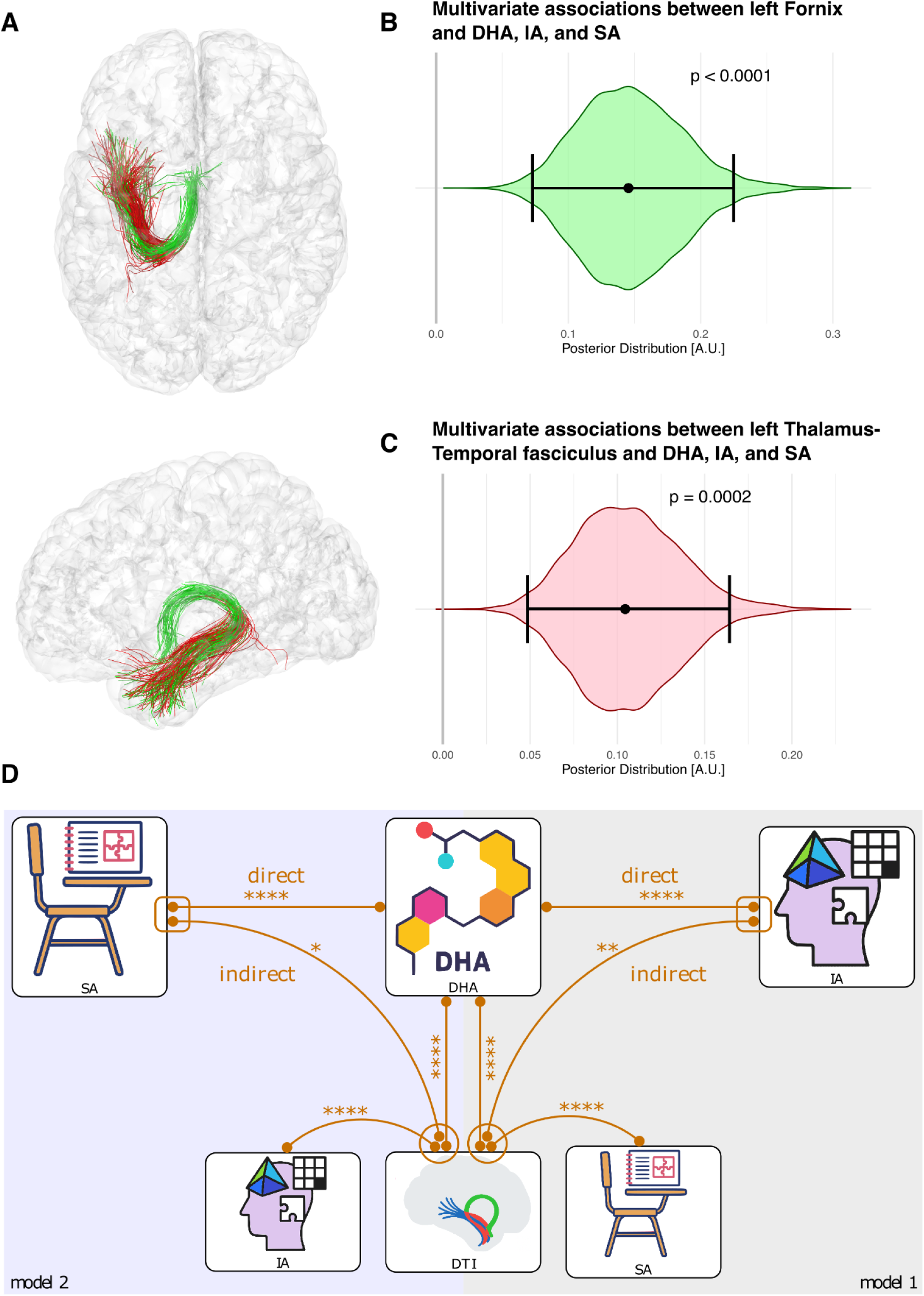
Associations between fractional anisotropy (FA) in deep white matter tracts and docosahexaenoic acid (DHA), school achievement (SA), and intellectual ability (IA). **A.** Tractography reconstructions of the left fornix (green) and the left thalamus–temporal fasciculus (red). **B–C.** Posterior distributions of the multivariate Bayesian LASSO associations for the left fornix (**B**, green) and the left thalamus–temporal fasciculus (**C**, red) with DHA, SA, and IA. Violin plots show posterior densities, with horizontal lines indicating medians and vertical lines representing the 95% highest density intervals. **D.** Summary of multivariate Bayesian LASSO models. Associations between DHA, SA, IA, and diffusion tensor imaging (DTI) metrics. Orange lines indicate statistically significant links (*p < 0.05, **p < 0.01, ***p < 0.001, ****p < 0.0001). Upper connections represent direct associations, whereas lower connections represent indirect associations via DTI. Rectangular nodes with outlined connectors denote variables jointly estimated within the same regression, while circles denote joint estimation in separate regressions. Results are presented for Model 1 (right) and Model 2 (left).

These findings motivated a follow-up multivariate analysis, conceptually similar to a mediation framework, to test whether IC5—or other brain components—contribute to an indirect pathway linking DHA to cognitive abilities. Because the correlation between DHA and SA/IA is strong, a conventional mediation analysis would have limited statistical power to identify an indirect pathway. Therefore, we implemented a multivariate Bayesian Lasso model with three dependent variables: DHA, SA, and IA. This joint modeling allowed us to identify brain components that simultaneously explained variance in two of the measures (e.g., DHA and SA) and then evaluate whether those same components predicted the third measure (e.g., IA) while controlling for DHA. In this framework, these steps are adjusted within the same model: the first contrast isolates the shared brain component between DHA and SA (or IA), and the second contrast tests whether this component predicts IA (or SA) beyond the direct effect of DHA. This approach was applied in two symmetric configurations: i) identifying the brain component jointly associated with DHA and SA, and testing its association with IA controlled by DHA; ii) identifying the brain component jointly associated with DHA and IA, and testing its association with SA controlled by DHA.

Both configurations showed that component IC5 was the strongest factor jointly explaining variance in DHA and SA (β_IC5_ median = –0.18, 95% HDI = [–0.29, –0.08], p_MV-LASSO-MCMC_ < 0.0001) or DHA and IA (β_IC5_ median = –0.17, 95% HDI = [–0.28, –0.07], p_MV-LASSO-MCMC_ < 0.0001).

In addition, the model identified two other significant components: IC13 (β_IC13_ median = –0.10, 95% HDI ≈ [–0.21, –0.006], p_MV-LASSO-MCMC_ = 0.02 (Model 1), 0.03 (Model 2)) and IC2 (β_IC2_ median = –0.07, 95% HDI ≈ [–0.15, –0.002], p_MV-LASSO-MCMC_ = 0.044 (Model 1), 0.046(Model 2)), which were consistent across both model configurations. Interestingly, in both configurations, the identified brain components were associated—independent of the direct DHA pathway—with IA or SA (see Figure 1D). Specifically, the shared brain component predicted IA when controlling for DHA (β_B→IA|DHA_ median = –0.59, 95% HDI = [–1.13, –0.14], p_MV-LASSO-MCMC_ = 0.008) and predicted SA when controlling for DHA (β_B→SA|DHA_ median = –0.33, 95% HDI = [–0.69, –0.01], p_MV-LASSO-MCMC_ = 0.029). Finally, to test whether our multivariate approach provides a deeper understanding of the data relationships, we assessed its fitting and predictability by comparing it with a null model that presented the same structure, but in which all beta variables were calculated independently in each of the three equations of the model. The Deviance Information Criterion (DIC) and Leave-One-Out Information Criterion (LOOIC) indicated that the multivariate model presented better data adjustment and data prediction, revealing that this approach better explains the data structure (ΔDIC = -35.9, ΔLOOIC = -24.9)

### Subcortical volumes, DHA, Scholastic Achievement, and Intellectual Ability

We then applied the same multivariate approach to investigate the relationship between subcortical volumes, DHA, SA, and IA, focusing on potential indirect pathways by which DHA may influence brain structure, which in turn impacts IA and SA. Using FreeSurfer subcortical segmentation, we excluded ventricular volumes and controlled for total intracranial volume, sex and socio-economical status. Our multivariate analyses identified four independent components (ICs) that captured the associations of interest, following the same model configurations applied in the FA analysis. Specifically, the IC5 (β_IC5_ median = 0.198, 95% HDI = 0.082–0.324, p_MV_LASSO_MCMC_ = 0.0006), the IC6 (β_IC6_ median = –0.195, 95% HDI = –0.308 to –0.086, p_MV_LASSO_MCMC_ = 0.0006), the IC7 (β_IC7_ median = 0.134, 95% HDI = 0.030–0.246, p_MV_LASSO_MCMC_ = 0.0078), and the IC13 (β_IC13_ median = –0.117, 95% HDI = –0.233 to –0.007, p_MV_LASSO_MCMC_ = 0.025). The shared brain component predicted IA when controlling for DHA (β_B→IA|DHA_ median = 0.40, 95% HDI = [0.007, 0.86], p_MV_LASSO_MCMC_ = 0.045) and predicted SA when controlling for DHA (β_B→SA|DHA_ median = 0.54, 95% HDI = [0.16, 0.98], p_MV_LASSO_MCMC_ = 0.003). Using all significant components and by multiplying the posterior distribution of the coefficient by the segmentations’ loadings that constitute each component, we observed positive associations for the volumes of the brain stem, bilateral pallidum, bilateral thalamus, and bilateral cerebellum

### Cortical areas volumes, DHA, intellectual ability, and scholastic achievement

Replicating the same approach, we analyzed gray matter volume and its associations with DHA, IA, and SA, modeling potential indirect effects of DHA on IA and SA via cortical morphology. Cortical volumes were estimated using FreeSurfer’s cortical segmentation based on the Desikan–Killiany atlas, and the analyses were controlled for total intracranial volume, sex and socio-economical status. Our multivariate analyses identified ICs that captured the associations of interest, following the same model configurations applied in the preceding analysis.

We found that five components significantly influence the association between gray matter volumes and DHA, IA, and SA (β_IC1_ median = -0.15, 95% HDI = -0.06 to –0.23, p_MV_LASSO_MCMC_ = 0.0006, β_IC4_ median = -0.18, 95% HDI = -0.29 to -0.06, p_MV_LASSO_MCMC_ = 0.002, β_IC7_ median = -0.14, 95% HDI = -0.26 to –0.03, p_MV_LASSO_MCMC_ = 0.0094, β_IC12_ median = -0.12, 95% HDI = -0.24 to –0.02, p_MV_LASSO_MCMC_ = 0.016, β_IC13_ median = 0.12, 95% HDI = 0.01 to 0.25, p_MV_LASSO_MCMC_ = 0.027). The shared brain component predicted SA when controlling for DHA (β_B→SA|DHA_ median = 0.32, 95% HDI = [0.1, 0.57], p_MV_LASSO_MCMC_ = 0.002) but did not predict IA when controlling for DHA (β_B→IA|DHA_ median = 0.23, 95% HDI = [-0.08, 0.57], p_MV_LASSO_MCMC_ = 0.14). Using all significant components and by multiplying the posterior distribution of the coefficient by the segmentations’ loadings that constitute each component, we observed positive associations for the volumes of gray matter that are illustrated in Figure 3.

**Figure 2.**
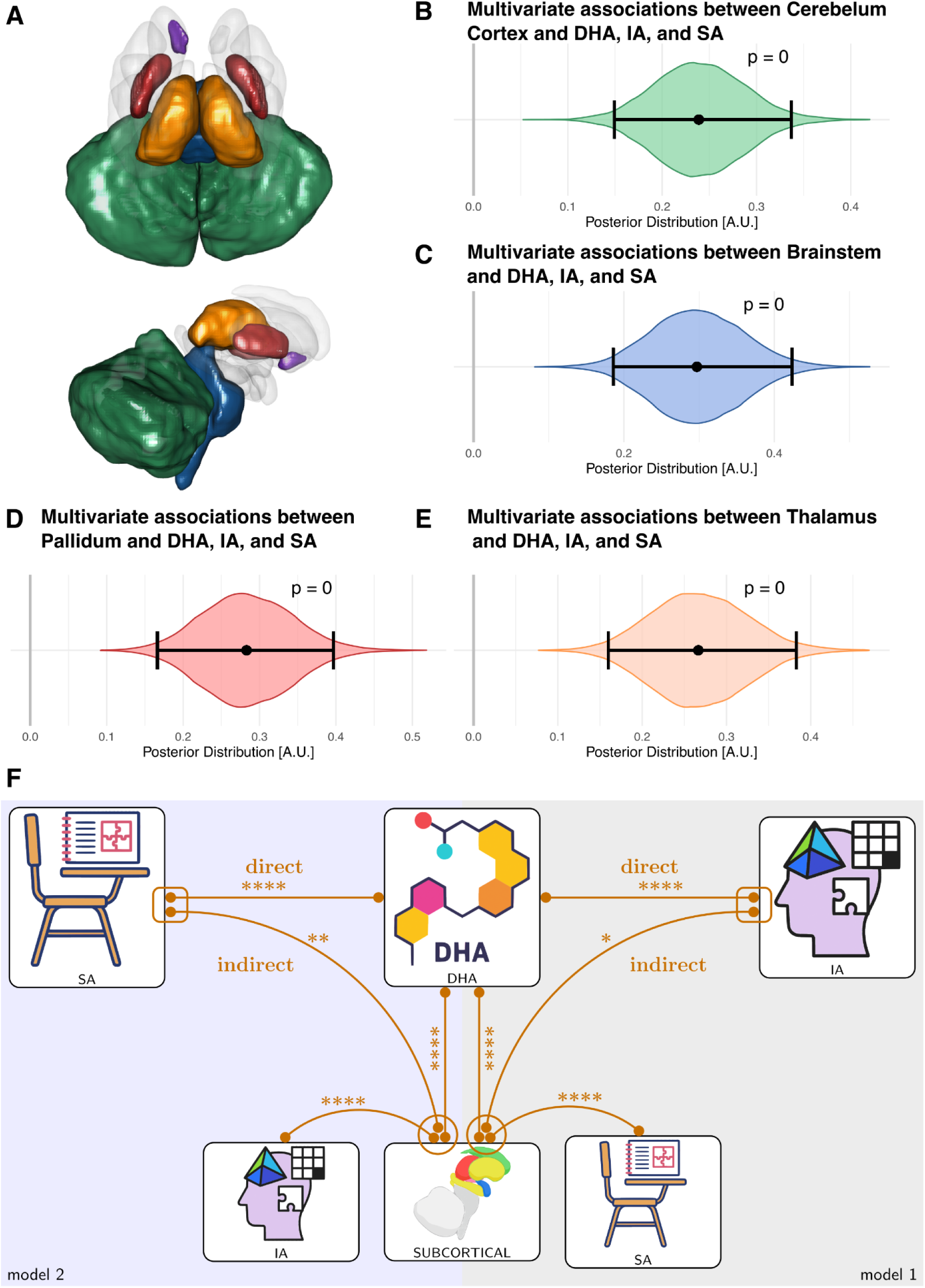
Associations between subcortical volumes and docosahexaenoic acid (DHA), school achievement (SA), and intellectual ability (IA). **A**. Segmented area included in the analysis, coloring area represents a significant area included in the IC detected in the multivariate analyses. **B-F.** Posterior distributions of the multivariate Bayesian LASSO associations for the right Cerebellar Cortex (**B**, green), the Brainstem (**C**, blue), the right Pallidum (**D**, red), and the right Thalamus (E, yellow) with DHA, IA, and SA. Violin plots show posterior densities, with horizontal lines indicating medians and vertical lines representing the 95% highest density intervals. Posterior distribution on the relación of the specific subcortical areas with IA, DHA, and SA. Summary of multivariate Bayesian LASSO models. Associations between DHA, SA, IA, and subcortical volumes. Orange lines indicate statistically significant links (*p < 0.05, **p < 0.01, ***p < 0.001, ****p < 0.0001). Upper connections represent direct associations, whereas lower connections represent indirect associations via DTI. Rectangular nodes with outlined connectors denote variables jointly estimated within the same regression, while circles denote joint estimation in separate regressions. Results are presented for Model 1 (right) and Model 2 (left).

**Figure 3.**
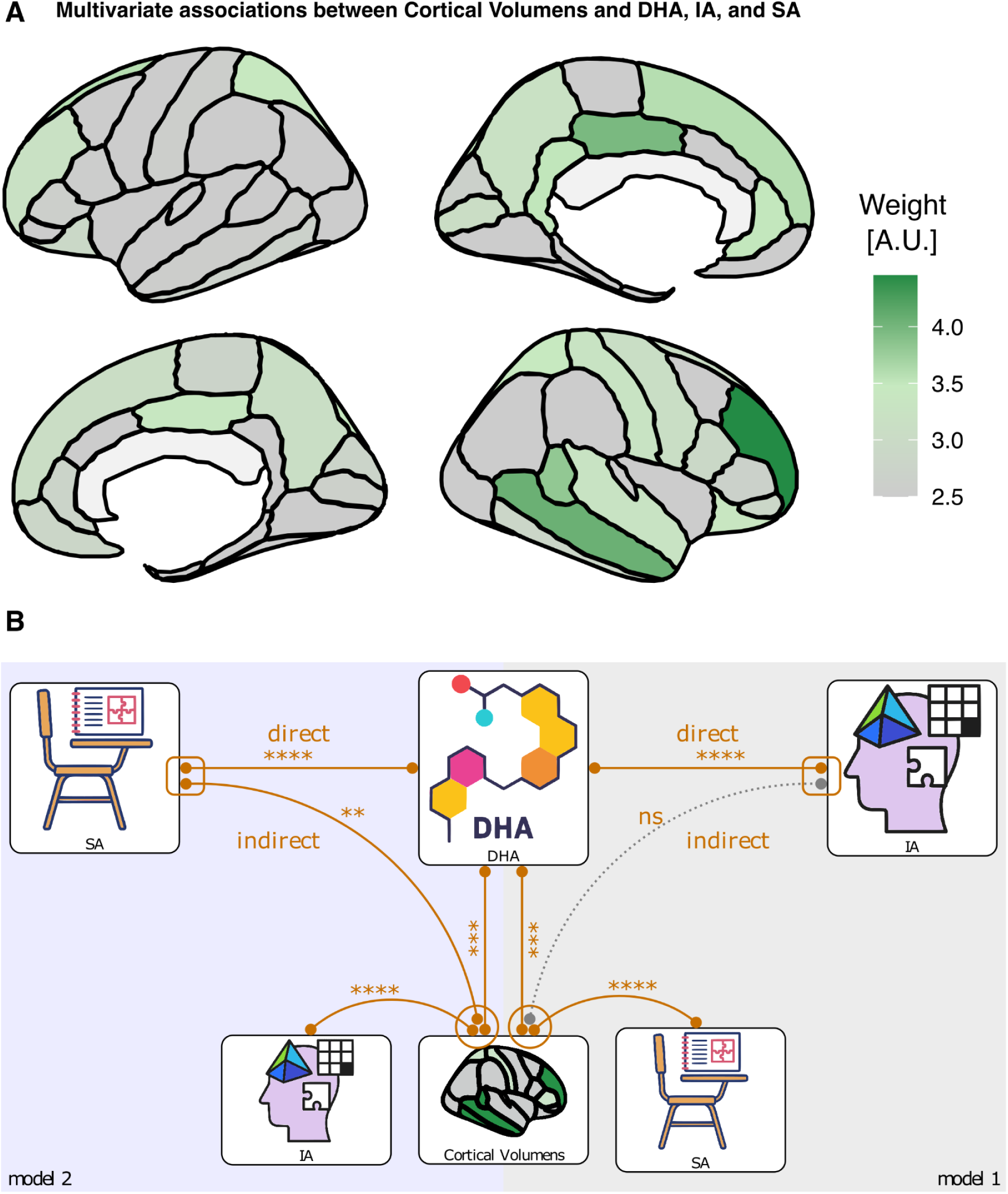
Associations between Cortical volumes and docosahexaenoic acid (DHA), school achievement (SA), and intelligence assessments (IA). **A**. Segmented area included in the analysis, coloring area represents a significant area included in the IC detected in the multivariate analyses. **B.** Summary of multivariate Bayesian LASSO models. Associations between DHA, SA, IA, and Cortical volumes. Orange lines indicate statistically significant links (*p < 0.05, **p < 0.01, ***p < 0.001, ****p < 0.0001). Upper connections represent direct associations, whereas lower connections represent indirect associations via cortical volumes.. Rectangular nodes with outlined connectors denote variables jointly estimated within the same regression, while circles denote joint estimation in separate regressions. Results are presented for Model 1 (right) and Model 2 (left).

**Figure 4.**
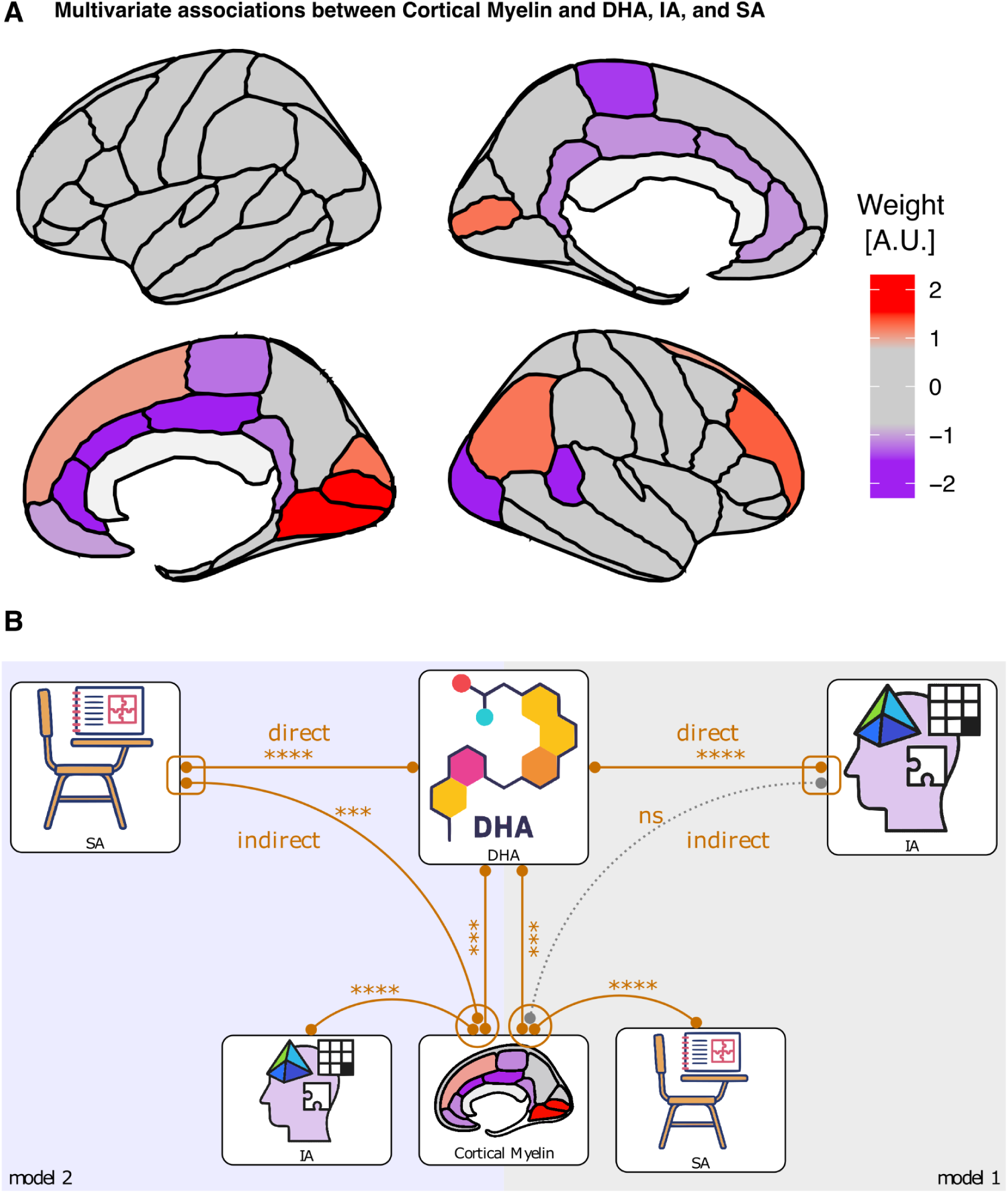
Associations between cortical myelin and docosahexaenoic acid (DHA), school achievement (SA), and intelligence assessments (IA). **A**. Segmented area included in the analysis, coloring area represents a significant area included in the IC detected in the multivariate analyses. **B.** Summary of multivariate Bayesian LASSO models. Associations between DHA, SA, IA, and Cortical myelin. Orange lines indicate statistically significant links (*p < 0.05, **p < 0.01, ***p < 0.001, ****p < 0.0001). Upper connections represent direct associations, whereas lower connections represent indirect associations via cortical myelin. Rectangular nodes with outlined connectors denote variables jointly estimated within the same regression, while circles denote joint estimation in separate regressions. Results are presented for Model 1 (right) and Model 2 (left).

**Figure 5.**
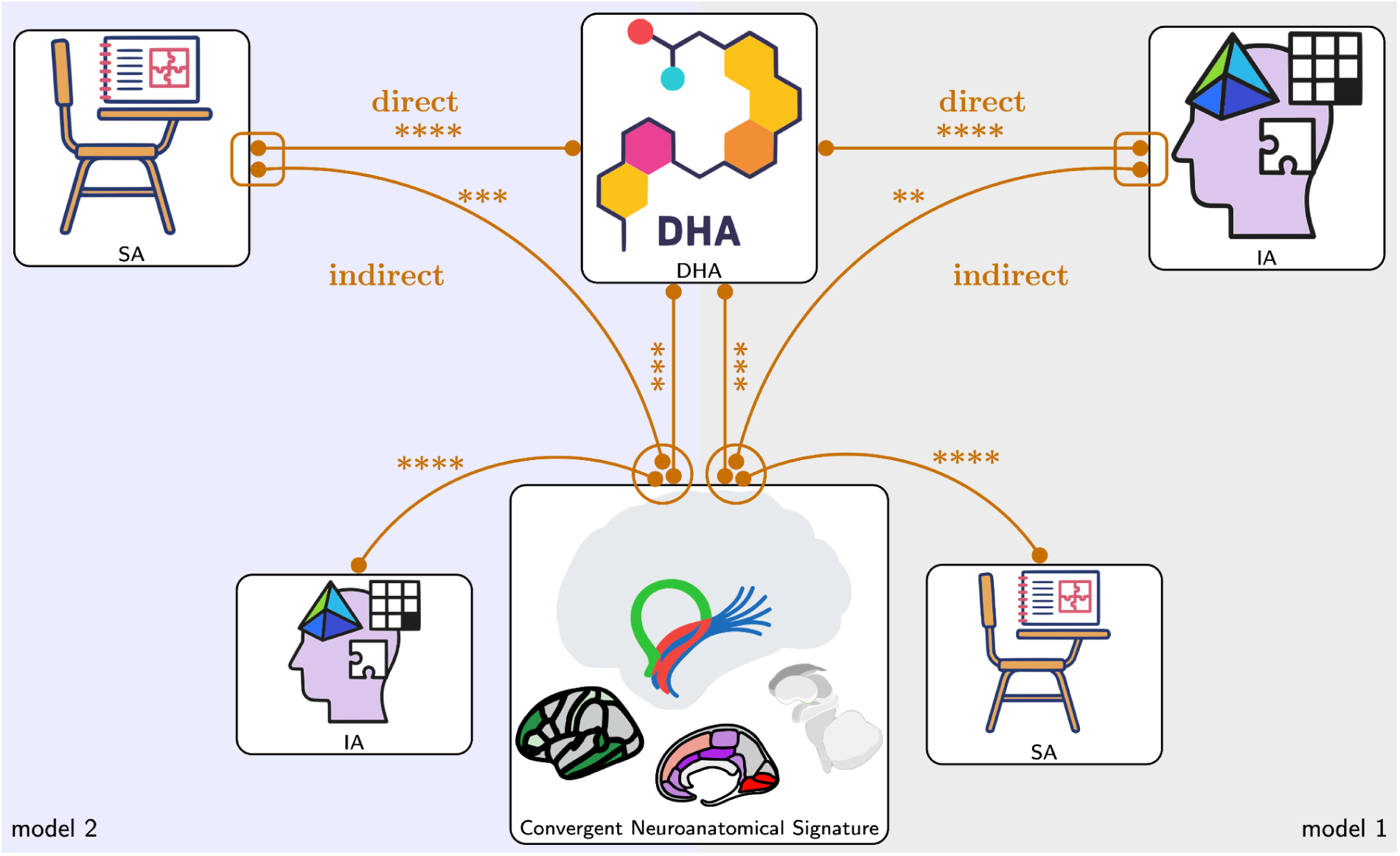
System Architecture: Direct and Indirect Pathways Linking DHA Status to Adolescent Cognitive Performance Across Multimodal Brain Measures. The diagram summarizes the final multivariate model integrating the most significant components across all four magnetic resonance imaging (MRI) modalities (deep white-matter microstructure FA, subcortical volumes, cortical volumes, and cortical myelin) to explain the association between erythrocyte docosahexaenoic acid (DHA) status, Scholastic Achievement (SA), and Intellectual Ability (IA). Central Nodes: The central panel (All Brain measures) represents the robust brain component that captures the variance jointly explained by DHA, SA, and IA. Direct Connections (Upper Lines): Lines connecting DHA directly to SA and IA illustrate the direct associations. Indirect/Mediation Pathways (Lower Lines): Lines connecting the combined brain component to SA and IA, while controlling for the direct effect of DHA, represent the indirect or mediation-like pathways. Statistical Significance: Orange lines denote statistically significant links based on the Multivariate Bayesian LASSO models (Models 1 and 2). The strength of the association is indicated by asterisks: (∗p<0.05, ∗∗p<0.01, ∗∗∗p<0.001, ∗∗∗∗p<0.0001).

**Figure 6.**
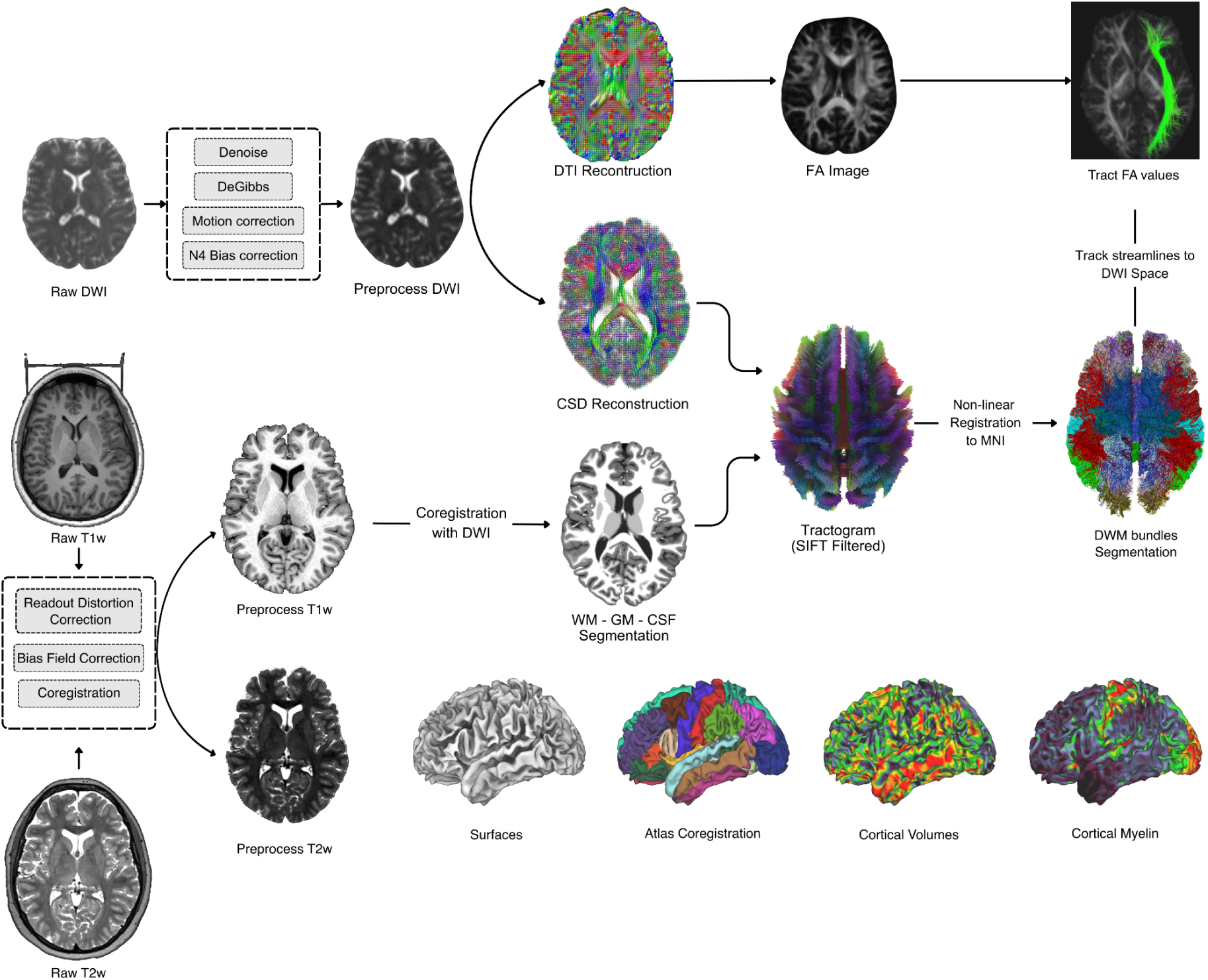
Imaging Processing Methodology. Raw diffusion-weighted images (DWI) undergo preprocessing steps including denoising, Gibbs ringing removal, motion correction, and N4 bias field correction. Following preprocessing, diffusion tensor imaging (DTI) reconstruction, fractional anisotropy (FA) maps, and constrained spherical deconvolution (CSD) tractography are computed. Probabilistic whole-brain tractography is generated and filtered using spherical-deconvolution informed filtering of tractograms (SIFT). Raw T1-weighted (T1w) and T2-weighted (T2w) images are preprocessed using the Human Connectome Project (HCP) pipeline, including readout distortion correction, bias field correction, and co-registration. The T1w, T2w, and DWI images are co-registered to a common space. White matter (WM), gray matter (GM), and cerebrospinal fluid (CSF) segmentation is performed following co-registration. Cortical surfaces are reconstructed, and cortical parcellation is generated using FreeSurfer’s SynthSeg algorithm based on the Desikan-Killiany atlas. Atlas co-registration enables extraction of cortical volumes. A structural connectome is constructed by identifying fiber bundles connecting specific regions of interest (ROIs) using the tractogram and cortical parcellation. Mean FA values of these tract-specific connections are calculated for statistical analysis. Additionally, T1w/T2w ratio maps are computed to estimate cortical myelin concentration.

### Cortical myelin, DHA, intellectual ability, and scholastic achievement

The last brain parameter extracted was the mean level of cortical myelin, obtained using a multivariate sparse approach to investigate its associations with DHA, IA, and SA. We modeled potential indirect pathways through which DHA may influence IA and SA via cortical morphology. Cortical myelin was estimated with the HCP pipeline, based on T1/T2-weighted images, and FreeSurfer’s cortical segmentation based on the Desikan–Killiany atlas, and all analyses were adjusted for total intracranial volume, sex and socio-economical status. We found that one component significantly modulated the associations between gray matter volumes and DHA, IA, and SA (β_IC2_ median = 0.23, 95% HDI = [0.13, 0.35], p_MV-LASSO-MCMC_ < 0.0001). The shared brain component predicted SA when controlling for DHA (β_B→SA|DHA_ median = 0.5, 95% HDI = [0.23, 0.83], p_MV-LASSO_MCMC_ = 0.0002) but did not predict IA when controlling for DHA (β_B→IA|DHA_ median = 0.31, 95% HDI = [-0.04, 0.69], p_MV_LASSO_MCMC_ = 0.07). Using all significant components and by multiplying the posterior distribution of the coefficient by the segmentations’ loadings that constitute each component, we observed positive and negative associations for the volumes of gray matter that are illustrated in Figure 3.

### Convergent neuroanatomical signature, DHA, intellectual ability, and scholastic achievement

Finally, we explored, within a unique multivariate model, the components that showed significant associations across the four brain measurements, with the goal of identifying the features most relevant to mediating the relationship between DHA and cognitive performance. Interestingly, four components reached statistical significance in this analysis. The strongest association was detected for FA through IC_5-FA_ (β_IC5–FA_ median = -0.14, 95% HDI = [-0.25, -0.05], p_MV-LASSO-MCMC_ = 0.0024). A component for cortical myelin also showed a significant effect (β_IC2–Mye_ median = 0.13, 95% HDI = [0.01, 0.25], p_MV-LASSO-MCMC_ = 0.018). Finally, two components for cortical gray matter volume were also statistically significant (β_IC1–GM_ median = -0.09, 95% HDI = [-0.17, -0.008], p_MV-LASSO-MCMC_ = 0.026; β_IC4–GM_ median = –0.10, 95% HDI = [-0.22, -0.002], p_MV-LASSO-MCMC_ = 0.045).

Consistent with prior analyses, the shared brain component predicted SA when controlling for DHA (β_B→SA|DHA_ median = 0.45, 95% HDI = [0.20, 0.71], p_MV-LASSO-MCMC_ = 0.0004) and predicted IA when controlling for DHA (β_B→IA|DHA_ median = 0.43, 95% HDI = [0.08, 0.76], p_MV-LASSO-MCMC_ = 0.005). The multivariate model provided a substantially better fit and predictive performance than the FA-only multivariate model (ΔDIC = –80.5, ΔLOOIC = –74.1), and also outperformed the null model (multiple univariate estimations of all brain features) (ΔDIC = –21.4, ΔLOOIC = –9.7). The findings remained significant after additionally controlling for EPA or AA levels (β_IC5–FA_ p_MV-LASSO-MCMC_ = 0.0006; β_B→SA|DHA-EPA_ p_MV-LASSO-MCMC_ < 0.0001), without evidence of improved model fit (ΔDIC = 3.4, ΔLOOIC = 1.5). Moreover, when EPA or AA was included as an additional dependent variable in a multivariate model, the overall pattern of results was unchanged (β_IC5–FA_ p_MV-LASSO-MCMC_ < 0.0001; β_B→SA|DHA-EPA_ p_MV-LASSO-MCMC_ < 0.0001), and EPA and AA showed no shared variance with brain structure or cognition (β_B→EPA_ p_MV-LASSO-MCMC_ = 0.99, β_B→AA_ p_MV-LASSO-MCMC_ = 0.14). These findings show that a shared multimodal component explains a meaningful portion of the DHA–cognition association, and that the multivariate approach better captures the coordinated variation across brain modalities specifically with DHA. This integrative signature offers insight into the neurobiological processes through which DHA may support cognitive performance.

## Discussion

In a Chilean community-based cohort of adolescents, we provide mechanistic evidence that individual differences in DHA status are coupled with cognitive performance through a system-level architecture of brain structure. Specifically, using multimodal MRI and mediation-inspired Bayesian Lasso models across deep white-matter microstructure, subcortical volumes, cortical volumes, and cortical myelin, we consistently identified a common deep white-matter component (including thalamus-temporal fasciculus and fornix) as the principal shared pathway linking higher erythrocyte DHA levels with superior cognitive abilities (measure may both SA and IA). This finding moves beyond established DHA-behavior correlations by pinpointing specific, modifiable neuroanatomical substrates that mediate cognitive development during the sensitive period of adolescence, with the deepest white-matter pathways showing the strongest contribution.

Additional brain components including cortical volumes ans cortical myelin also contributed significantly revealing a system-component also relevenat in cogntive performabe. Tract-level loadings highlighted positive contributions from the left fornix and the left thalamus–temporal fasciculus as key node of the brian features. Joint models that controlled for the direct effect of DHA (and other fatty acids like EPA and ALA) showed that these brain components predicted the complementary cognitive outcome (IA given SA, and SA given IA), supporting a mediation-like architecture in which DHA may influence cognition both directly and indirectly through its association with structural brain networks.

Adolescence is a period of rapid remodeling of long-range connectivity, myelination, and subcortical–cortical integration [40]. Our results are consistent with the view that DHA—an omega-3 fatty acid enriched in neuronal membranes—relates to this remodeling in ways that bear on cognitive function [41–44]. The fornix and thalamus–temporal fasciculus, which carried the dominant tract loadings, are well positioned to influence learning and reasoning: the fornix is the principal efferent pathway of the hippocampal formation, supporting memory-guided inference and knowledge consolidation [45], whereas the thalamotemporal system contributes to cortico-cortical communication and multimodal integration [46–48]. This track seems to underlie the support for cortical associations that have been described in several cognitive abilities [49,50], including cognitive control, working memory, and decision making [51–60]. Our findings provide convergent associations linking these tracts with DHA, SA, and IA, thereby offering a plausible substrate for the observed behavioral effects.

Cortical myelin and cortical volumes also contributed unique variance, with a myelin component predicting SA (beyond DHA) more robustly than IA. This asymmetry suggests that school performance—as a composite of sustained engagement, reading/maths skills, and executive routines—may be especially sensitive to myelin-related features that optimize conduction velocity and temporal coordination across networks [22,61–64]. By contrast, IA may rely more on distributed deep-white-matter and subcortical architecture captured by the dominant white-matter component [65–69] subcortical (brainstem [70,71], thalamus [48], pallidum [72], cerebellum [73]) and cortical[74] contributions align with roles in arousal control, relay/attention, action selection, and supervised learning, respectively, and reinforce the interpretation of DHA effects as presenting a widespread and a focal effect component.

At the component level, the key white-matter component showed negative regression coefficients, yet tract-resolved reconstructions revealed positive associations for the fornix and thalamus–temporal fasciculus. This reflects the indeterminacy of ICA signs and the fact that each component aggregates positive and negative tract loadings. Our posterior-weighted back-projection clarifies that the DHA–cognition pathway is driven by increased FA in specific tracts. The single-model multivariate LASSO further strengthens inference [50,75,76] by identifying brain components jointly associated with DHA and one cognitive outcome and then testing their predictive value for the other outcome while controlling for DHA, we reduce the risk that associations reflect parallel but unrelated correlates of DHA. Although not a formal causal mediation test, the convergent pattern across two symmetric configurations and four modalities argues for an indirect pathway consistent with the mediation model [77,78].

Multiple biological routes could link DHA status to the observed macrostructural signatures. DHA modulates membrane fluidity, receptor and channel function, synaptic vesicle dynamics, and neuroinflammatory tone [2,79,80]; it is also a critical constituent of myelin lipids [81,82]. In adolescents, such cellular effects may scale to measurable differences in axonal packing/coherence (reflected in FA), regional volumes, and myelination indices derived from T1/T2 [20,22,81–84]. The prominence of thalamocortical and hippocampal pathways is notable given their protracted developmental trajectories and their sensitivity to metabolic and nutritional constraints [22]. Subcortical associations in brainstem and cerebellum may index DHA-related support of sensorimotor integration, timing, and vigilance—functions that scaffold both domain-general reasoning (IA) and applied academic performance (SA) [85].

Our findings replicate the positive association between erythrocyte DHA and cognitive outcomes previously reported in the broader cohort from which this neuroimaging subsample was drawn, and extend that work by identifying distributed neuroanatomical correlates that behave as shared pathways to both SA and IA [37]. Prior heterogeneous results in the literature may partly reflect reliance on dietary questionnaires rather than blood biomarkers [86,87], focus on single brain metrics [88], or samples with restricted nutritional variance [89]). By leveraging multimodal MRI and a unified multivariate framework, we show that DHA’s behavioral associations are best understood at the systems level: a backbone deep-white-matter component, supported by subcortical structure and cortical microstructure, collectively links nutritional status with cognition.

Methodologically, three features increase confidence in the results. First, the use of erythrocyte DHA provides a stable biomarker of status [12,90,91]. Second, the multimodal design (DTI, morphometry, myelin mapping) allows converging evidence across partially independent biological contrasts [92,93]. Third, the Bayesian multivariate LASSO—combined with ICA for dimensionality reduction and posterior back-projection for anatomical interpretation—offers principled regularization [94,95], joint modeling of coupled outcomes (DHA, SA, IA) [96], and propagation of uncertainty from component to tract level [97]. This Bayesian framework naturally quantifies uncertainty in high-dimensional neuroimaging settings [97], while ICA effectively reduces data dimensionality by extracting biologically meaningful spatial components that correspond to white matter tracts and functional networks [98,99].

Several constraints temper causal interpretation. The design is cross-sectional; thus, we cannot infer temporal precedence of DHA → brain → cognition [100]. Although we adjusted models and examined shared pathways, residual confounding (e.g., unmeasured dietary patterns, micronutrients, sleep, physical activity, or other environmental factors) could contribute [101–103]. Additionally, the study did not use standardized questionnaires to screen for undiagnosed neurodevelopmental conditions, such as autism spectrum disorder (ASD) or attention-deficit/hyperactivity disorder (ADHD). These conditions are associated with distinct structural brain differences, including variations in cortical thickness, gray and white matter volumes, and altered connectivity patterns [104,105]. FA is a composite metric influenced by myelination, axonal geometry, and crossing fibers; future work should incorporate multi-compartment models (e.g., NODDI, fixel-based analyses) and myelin-sensitive sequences to refine microstructural inferences [21,106,107]. Myelin estimates derived from T1/T2 ratios are indirect and can be affected by water content and iron; quantitative MRI (e.g., Magnetization Transfer Saturation [108], longitudinal relaxation rate [109], Effective transverse relaxation rate) would improve specificity [110,111]. Finally, generalizability beyond Chilean adolescents requires replication in cohorts with different nutritional environments and educational systems.

The present results highlight DHA sufficiency as a biologically plausible and potentially modifiable factor related to adolescent brain architecture and cognition [37]. Longitudinal and interventional studies [112] are now warranted to formally test mediation—ideally combining randomized DHA supplementation [113], repeated multimodal imaging [22,114], fine-grained educational outcomes, or the use of non-invasive brain stimulation to address the underlying neurobiological pathway [115]. Inclusion of mechanistic biomarkers, genotypes relevant to fatty-acid metabolism [116], and objective measures of sleep and physical activity would help disentangle pathways [117,118]. Network-level analyses (effective connectivity, structural–functional coupling [49]) may clarify how thalamohippocampal and cerebello-thalamo-cortical loops translate nutritional status into academic and reasoning performance. From a public health perspective, triangulating biomarker-based nutritional surveillance with brain metrics in school-based programs could inform policies aimed at supporting cognitive development during this sensitive window [84,119].

DHA status relates to adolescent cognition through widespread neuroanatomical features, with a deep white-matter component—anchored by fornical and thalamotemporal pathways—emerging as the principal shared pathway and complemented by subcortical structure, cortical morphology, and cortical myelin. These findings move the field beyond simple DHA–behavior correlations, toward a systems neuroscience account in which nutritional lipids shape the architecture of developing brain networks that support both scholastic achievement and general intellectual ability.

## Methods

### Design

This is an observational, cross-sectional, and correlational study.

### Description of the population

As previously described [36–39], participants were drawn from a representative cohort of school-age students who had completed the University Selection Test (PSU). Briefly, a representative sample of 671 young people who graduated from high school in 2013 took the PSU, with 550 and 548 completing the language scholastic achievement (LSA) and mathematics scholastic achievement (MSA) tests, respectively. Only participants with high (n = 91) and low (n = 69) scholastic achievement (SA) in both tests were considered for the study. A total of 122 participants agreed to participate: 70 with high PSU outcomes (males n = 48) and 52 with low PSU outcomes (males n = 23). Finally, 99 participants were successfully scanned with T1- and T2-weighted imaging, and, of them, 93 additionally underwent diffusion-weighted imaging. This represents 69% and 56% of students with the highest and lowest PSU scores from the original selected sample, respectively.

### Intellectual ability (IA)

IA was assessed with the standard version of the Raven’s Progressive Matrices Test (RPMT) in book form, with a general scale for children of 12 years and above that had been standardized for Chilean school age students [120]. The Standard RPMT is a non-verbal test and, in any of its forms, constitutes one of the tests most frequently applied for quantification of general intelligence, evidencing a robust and reliable measure of the general intelligence factor [121]. The test was administered collectively in the classrooms by an educational psychologist. WHO experts for developing countries have recommended applying Raven’s test because its results are not affected by culture [122]. Scores were registered as a percentile scale according to age, in the following grading: Grade I = Superior Intellectual Ability (score ≥ p95) to Grade V = Intellectually Deficient (score ≤ p5).

### Scholastic achievement (SA)

SA was measured by means of the university selection test (“Prueba de Selección Universitaria”, PSU). Results from the PSU outcomes in language and mathematics tests were registered for the 2010 first HSG school-age students when they graduated from the fourth HSG in 2013. PSU has a maximum score of 850 and a minimum of 150 for each test (language and mathematics tests with 80 items each) and was expressed as mean ± SD. Scores below 450 bar students from applying to universities.

### Fatty acid profile analysis

Blood samples were taken from fasted male and female young for erythrocyte fatty acid assessment. The samples were immediately centrifuged to obtain the erythrocyte fraction (3000× g for 10 min at 20 °C), then butylated hydroxytoluene (BHT) was added to the blood samples as antioxidant and and then frozen at −80 °C until further analysis. Quantitative extraction of total lipids from erythrocytes was carried out according to Bligh and Dyer [123]. Erythrocytes samples were separately mixed with ice-cold chloroform/methanol (2:1 v/v), magnesium chloride was added (0.5 N), and the mixture was homogenized in an Ultraturrax homogenizer (Janke & Kunkel, Stufen, Germany). The total lipids extracted from erythrocytes were separated by thin layer chromatography (TLC) (aluminum sheets 20 × 20 cm, silica gel 60 F-254; Merck), using the solvent system hexane/diethylether/acetic acid (80:20:1 v/v). After the development of the plates and solvent evaporation, lipid spots were visualized by exposing the plates to a Camag UV (250 nm) lamp designed for TLC. The solvent system allowed the separation of phospholipids, triacylglycerols, cholesterol and cholesterol esters according to their relative mobility. Spots corresponding to phospholipids were scraped from TLC plates and extracted by elution with either diethyl ether or chloroform/methanol (2:1 v/v), according to Ruiz-Gutierrez et al. [124]. Fatty acid methyl esters (FAMEs) from erythrocyte phospholipids were prepared according to Morrison and Smith [125]. Briefly, samples had previously been dissolved in chloroform/methanol (2:1 v/v) and were then evaporated under nitrogen stream until the volume was halved, then boron trifluoride (12% methanolic solution) and sodium hydroxide (0.5 N methanolic solution) at a temperature between 90 and 95°C were added, then the mixture was cooled. FAMEs were extracted with 0.5 mL of hexane. FAMEs were identified and quantified by gas chromatography in an Agilent equipment (model 7890B, Santa Clara, CA, USA) equipped with a capillary column (Agilent HP-88, 100 m × 0.250 mm; I.D. 0.25 µm) and flame ionization detector (FID). The injector temperature was set at 250 °C and the FID temperature at 300 °C. The oven temperature at sample injection was initially set at 120 °C and was programmed to increase to 220 °C at a rate of 5 °C per min. Hydrogen was utilized as the carrier gas at a flow rate of 35 cm per second in the column, and the inlet split ratio was set at 20:1. Identification and quantification of FAMEs were achieved by comparing the retention times and the peak area% values of unknown samples to those of commercial lipid standard (Nu-Chek Prep Inc., Elysian, MN, USA). C23:0 was used as an internal standard (Nu-Chek Prep Inc., Elysian, MN, USA) and data was processed using the Hewlett-Packard Chemstation software system.

### Brain structural data acquisition

Images were acquired at the Radiology Department of the Clínica Alemana de Santiago with a 3 T Siemens Skyra scanner and a 20-channel head coil. All participants underwent (i) a sagittal 3D anatomical MPRAGE T1-weighted imaging (repetition time [TR]/ echo time [TE] = 2530/2.19 ms, inversion time [TI] = 1100 ms, flip angle = 7°; 1 × 1 × 1 mm3 voxels), (ii) a sagittal 3D anatomical SPC T2-weighted (TR/TE = 3200/412 ms, flip angle = 120°; echo train length [ETL] = 258; 1 × 1x1 mm3 voxels) and (iii) an axial 3D echo-planar imaging (EPI) (TR/TE = 8600/95 ms, 2 × 2 × 2 mm3 voxels, flip angle = 90°) with diffusion gradients applied in 30 non-collinear directions and two optimized b factors (b1 = 0 and b2 = 1000 s/mm2) with three repetitions. All imaging was acquired on a 3 T Siemens Skyra (Siemens AG, Erlangen, Germany) MR scanner with a gradient of 45 mT/m and a maximum slew rate of 200 mT/m/s.

### Imaging Processing

#### Structural MRI Analysis

T1-weighted and T2-weighted structural images were preprocessed using the Human Connectome Project (HCP) minimal preprocessing pipeline[126] and described in our prior work [49,50,127]. Briefly, The pipeline included gradient nonlinearity correction, readout distortion correction using spin-echo field maps, anterior commissure-posterior commissure (AC-PC) alignment, and bias field correction using the square root of the product of T1w and T2w images[126]. T1w and T2w images were rigidly aligned and used to generate T1w/T2w ratio maps for cortical myelin content estimation [128]. Cortical surface reconstruction and subcortical segmentation were performed using FreeSurfer 7.4 [129] integrated within the HCP pipeline. Cortical parcellation was generated using the SynthSeg algorithm [130], a robust deep learning-based segmentation method, based on the Desikan-Killiany atlas[131]. White matter, gray matter, and cerebrospinal fluid tissue segmentation were computed for anatomically constrained tractography. The resulting cortical surfaces, parcellations, and volumetric measures were used for structural connectome construction and cortical metric extraction.

#### Connectivity MRI Analysis

The diffusion imaging was processed using MRtrix3 software [132] The first stage of the process was to apply a pre-processing pipeline, which involved several steps designed to enhance image quality and reduce the impact of artifacts. These steps included image denoising, correction of Gibbs ringing, motion and eddy current distortion correction, and N4 bias field correction. The pre-processed diffusion images were used to calculate the Diffusion Tensor Imaging (DTI) model and extract the fractional anisotropy (FA). Preprocessed diffusion data were used to compute fiber orientation distributions using 3-tissue response function estimation and multishell multi-tissue Constrained Spherical Deconvolution [133]. Whole-brain probabilistic tractography was performed using the iFOD2 algorithm with Anatomically Constrained Tractography (ACT) [134] (step size 2mm x 2mm x 2mm, max degree = 45°, min length = 40, max length = 250) generating 10 million streamlines per subject. These were subsequently filtered using the Spherical deconvolution Informed Filtering of Tractograms (SIFT) [135] algorithm to 3 million streamlines per subject to reduce reconstruction biases and improve biological accuracy. T1w and dMRI images were co-registered using FSL software [136]. The T1w images were then normalized to MNI space using FSL non-linear transformation, and the tractography was transformed into MNI space. An automated deep white matter segmentation algorithm based on Euclidean distances[137] was employed to extract the 36 bundles using an DWM atlas [138]. The streamline bundles were tracked back to their original space where fractional anisotropy (FA) masks were computed for each bundle and subject to analyze intersubject variability.

### Statistical analysis

#### Independent component analysis (ICA)

Tract-wise FA values, Cortical Myline, Subcortical and Cortical Volumes were first robustly standardized by centering each tract on its median and scaling by its median absolute deviation (MAD). A data matrix X (subjects × tracts or Cortical or subcortical areas) was submitted to Independent Component Analysis (ICA) to identify independent modes of covariance. To obtain stable and reproducible components, the *fastICA* algorithm (R package *fastICA*) was run repeatedly (n_runs_=50) extracting n_comp_=30 components per run. For each run the mixing matrix A (tracts × components) was transposed so that rows corresponded to tracts and columns to components, and these matrices were collected across runs. To align components across runs, each run was matched to a reference run (the first iteration) by maximizing the absolute correlation between component maps. The linear assignment problem was solved using the Hungarian algorithm implementation (*solve_LSAP*, *clue* package), which returned a permutation aligning target components to the reference. Component sign ambiguities were resolved by multiplying each column by the sign of its correlation with the corresponding reference column. The aligned mixing matrices were then averaged across runs to produce a stable group-level mixing matrix Ā. Subject-level component scores S were obtained by projecting the standardized data onto the pseudoinverse of the averaged mixing matrix: W=pinv(Ā) and S = XW^T^. Finally, components were reordered by the amount of variance explained across subjects (i.e., components were sorted in decreasing order of variance of the columns of S), and the final Ā and S matrices were saved for subsequent analyses.

#### Bayesian LASSO

Subject-wise ICA scores (S) served as predictors in Bayesian regression models to relate brain-derived components to behavioral and cognitive outcomes. A Bayesian LASSO specification was coded in JAGS (via *rjags*/*runjags*) following XX. The likelihood assumed a Gaussian distribution for each outcome y with mean μ_i_=β_0_+∑^p^_j=1_ β_j_X_ij_ and precision parameter 𝜏 = 1 / σ^2^. Weakly informative normal priors were assigned to intercepts (e.g., β0∼𝑁(0, 10^2^) and gamma priors to precision parameters. Adaptive LASSO-style shrinkage was imposed by defining hierarchical priors on regression coefficients: β_j_∼N(0,𝜏_βj_−1) with 𝜏_𝛽𝑗_=1/(𝜎^2^ 𝜏_𝑗_^2^) and 𝜏_𝑗_^2^∼Exp(𝜆^2^/2). The global penalty parameter 𝜆 was computed from the data following the heuristic implemented: 𝜆 = p×Var(residuals_OLS_) / ∑β_OLS_, where β_OLS_ are coefficients from an ordinary least squares fit of y ∼ X_j_. Subject-level covariates (e.g., standardized socioeconomic index and sex) were appended to the predictor matrix when specified. MCMC sampling was performed with three parallel chains. Chains were initialized with deterministic starting values and distinct JAGS RNG seeds to ensure reproducible sampling. Models were adapted for 1,000 iterations, followed by a burn-in of 5,000 iterations. Posterior samples were then drawn with extensive iterations (script defaults produce tens of thousands of iterations per chain with thinning; stored posterior samples depend on thinning and total iterations), and posterior chains from all three chains were concatenated for inference. Convergence was assessed using the Gelman–Rubin potential scale reduction factor (R^) and Geweke diagnostics; R^ values < 1.1 were taken as evidence of adequate between-chain convergence. Posterior summaries include medians and 95% highest-density intervals (HDIs) for parameters of interest. A Bayesian (two-tailed) *p*-value was computed as twice the smaller of the proportions of posterior samples above or below zero, i.e. p_MCMC_= 2 min(P(β>0), P(β<0)).

#### Component-to-measure contribution and visualization

For interpretation of significant component–behavior associations, measure-level contributions (track, cortical or subcortical areas) were derived by combining the averaged mixing matrix Ā and posterior coefficient estimates. For each component identified as significant, its tract weights (rows of Ā) were multiplied by the corresponding posterior coefficient (or a standardized summary of that coefficient) and summed across significant components to yield an aggregate tract-level score. Posterior distributions of these tract-weighted summaries were inspected and visualized using violin plots and ordered bar plots to illustrate each tract’s relative contribution and uncertainty. Reported inference therefore reflects the full posterior distribution propagated from component weights and regression coefficients.

### Multivariate regressions

#### Step 1: Brain signature of DHA

In the first stage, we modeled DHA as a function of first independent components from each of the four brain measures that accumulated the 90 % of the variance (FA from deep tracts, volumes of subcortical areas, volumes of cortical parcellations, and myelin of cortical parcellations) using a Bayesian LASSO prior for coefficient shrinkage and variable selection indicated in the preceding section. This model is expressed by the following equation:

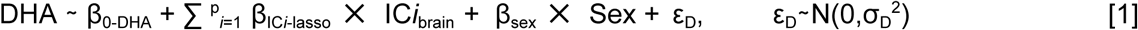

This approach yields a sparse neuroanatomical profile of association with DHA, identifying the most salient and predictive components of brain structure. For each participant, we then computed an individualized DHA Brain Signature Score. This score was calculated as the sum of product of the participant’s brain data and the model’s sparse weight vector:

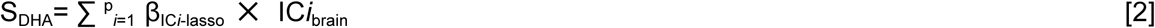

#### Step 2: Multivariate regressions for the Brain signature associated with DHA, SA, and IA

For the multivariate analysis, we fitted three regression models within a joint modeling framework. This approach allowed the S_DHA_ component to capture not only the associations between DHA and specific brain measures but also the shared neuroanatomical profile jointly related to DHA, scholastic achievement (SA), and intellectual ability (IA). To test indirect pathways, we implemented two symmetric models. In Model 1, the regression for IA included the direct pathway from DHA, whereas in Model 2, the regression for SA included this direct pathway. In this configuration, the association between S_DHA_ and the cognitive measures can be interpreted as an indirect pathway: DHA influences brain structure, which in turn impacts IA or SA, beyond the direct effect of DHA. Specifically, we estimated the following equations (in addition to Equation 1):

Model 1

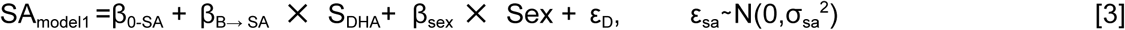

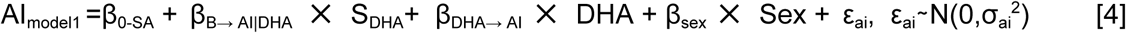

Model 2

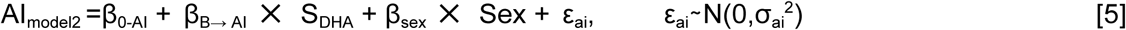

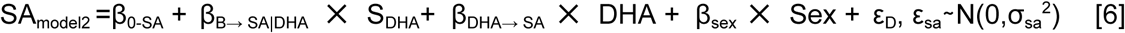

### Ethical approval and consent to participate

The experimental protocol and all methods were performed in accordance with institutional guidelines and were approved by the Ethics Committee in Studies in Humans of the Institute of Nutrition and Food Technology Dr. Fernando Monckeberg Barros (INTA), University of Chile, and ratified by the Bioethics Committee of the National Fund for Scientific and Technological Development (FONDECYT), Chile. The participants’ informed consent was obtained according to the norms for Human Experimentation, Code of Ethics of the World Medical Association (Declaration of Helsinki).

## Foundings

This work was supported by Grants 1100431 (DI), 1150524 (DI) and 1251073 (PB) from the National Fund for Scientific and Technological Development (FONDECYT). The funders had no role in study design, data collection, and analysis, decision to publish, or preparation of the manuscript.

## Acknowledgements

The authors are grateful to the Agency for Quality Education, the Studies Center of the Ministry of Education of Chile, and the Department of Evaluation, Measurement, and Educational Registry (DEMRE) of the University of Chile for the facilities to carry out this research, which reviewed, approved, and authorized this study, and for providing PSU outcomes.

## Author contributions

D.I., P.B., A.F-V, R. V., F.Z. developed the conceptualization of the study and analysis; D.I., A.A., R.V., and C.B. participated in data collection; C.L. and C.S. performed the analysis for brain development study MRI exams; P.B., A.F-V, P.G., F.Z., and S.N., performed the analysis for brain parameters; D.I., P.B., and A.F-V planned the application of statistical, mathematical and computational techniques to analyze or synthesize study data, and prepared figures; D.I. and V.A. applied and analyzed IA tests; D.I. was the principal investigator for funding acquisition; D.I., P.B., A.F-V, P.S-I. wrote the original draft. All authors reviewed and approved the final version to be published and agreed to be accountable for the integrity and accuracy of all aspects of the work.

